# Structural and functional insights into the mechanism of action of plant boron transporters

**DOI:** 10.1101/2020.12.15.422910

**Authors:** Savvas Saouros, Thotegowdanapalya C. Mohan, Cristina Cecchetti, Silke Lehmann, Joseph D. Barritt, Nicola J. Scull, Paul Simpson, Yilmaz Alguel, Alexander D. Cameron, Alexandra M. E. Jones, Bernadette Byrne

**Affiliations:** Department of Life Sciences, Imperial College London, Exhibition Road, London, SW7 2AZ; School of Life Sciences, University of Warwick, Gibbet Hill Road, Coventry CV4 7AL, UK

## Abstract

Boron has essential roles in plant growth and development. BOR proteins are key in the active uptake and distribution of boron, and regulation of intracellular boron concentrations. However, their mechanism of action remains poorly studied. BOR proteins are members of the SLC4 family of transporters and thus homologues of well studied mammalian transporters including the human Anion Exchanger 1 (hAE1). Here we generated *Arabidopsis thaliana* BOR1 (AtBOR1) variants based i) on known disease causing mutations of hAE1 (S466R, A500R) and ii) a loss of function mutation (D311A) identified in the yeast BOR protein, ScBOR1p. The AtBOR1 variants express in yeast and localise to the plasma membrane, although both S466R and A500R exhibit lower expression than the WT AtBOR1 and D311A. The D311A, S466R and A500R mutations result in a loss of boron efflux activity in a yeast *bor1p* knockout strain. *A. thaliana* plants containing these three individual mutations exhibit substantially decreased growth phenotypes in soil under conditions of low boron. These data confirm an important role for D311 in the function of the protein and show that mutations equivalent to disease causing mutations in hAE1 have major effects in AtBOR1. We also obtained a low resolution cryo-EM structure of a BOR protein from *Oryza sativa*, OsBOR3 lacking the 30 C-terminal amino acids. This structure confirms the gate and core domain organisation previously observed for related proteins, and is strongly suggestive of an inward facing conformation.

## Introduction

Boron (B) is a chemical element key for plant growth and development, playing essential roles in formation and stability of the plant cell wall and is crucial for normal leaf expansion, through the cross-linking of the polysaccharide rhamnogalacturonan II (RG-II) (Hu and Brown, 1994). Borate-diol diester bridges form between the apiosyl residues of two monomers of RG-II (Chormova and Fry, 2015). Boron deficiency or excess causes a range of plant growth defects (Dell and Huang, 1997; Li et al., 2001; Camacho-Cristóbal et al., 2015) and is a major issue for crop growth (Tanaka and Fujiwara, 2008). The optimal B concentration is narrow in many plants; in wheat it is between 10 μg/g and 100 μg/g tissue concentration (Shorrocks, 1997). B is highly soluble and thus is readily leached from the soil in areas with high rainfall including parts of the USA and China (Miwa and Fujiwara, 2010). The resulting B deficiency limits plant growth, root elongation, fruit ripening and seed production (Goldbach et al., 2001). Conversely excess B is often the result of low rainfall in areas of, for example, Australia, Turkey and South America and the Middle East and reduces root cell division and chlorophyll content, which in turn limits growth of shoots and roots (Pallotta et al., 2014).

Plants have a complex system of channels and transporters involved in the uptake and distribution of Boron (Hrmova et al., 2020). Boron is taken up from the soil in the form of boric acid, B(OH_3_), via a passive process, through a combination of direct diffusion across the membrane and facilitated diffusion through the nodulin 26-like intrinsic proteins (NIPs) (Takano et al., 2006). Borate transporters (BORs) are responsible for active efflux of boron from cells, most likely in the charged borate [B(OH_4_)]^−^ form (Miwa and Fujiwara, 2010). The BORs are thus suggested to be borate antiporters coupling transport of the substrate to the H^+^ gradient (Jennings et al., 2007). The first identified and best characterised BOR protein, BOR1 from *Arabidopsis thaliana* (AtBOR1), is responsible for active export of B from the root cells to the xylem for further distribution around the plant (Takano et al., 2002). Overexpression of AtBOR1 leads to more efficient uptake of B and resistance to B deficiency (Uragachi et al., 2014). BOR proteins are also key in preventing toxic accumulation of B in cells. Increased levels of AtBOR4 from *A. thaliana* (Miwa et al., 2007) and the homologue Bot1 from barley (Sutton et al., 2007) confer increased tolerance of high B concentrations by exporting the excess B back into the soil. AtBOR1 is regulated at least in part by cellular B concentration, with low B concentration resulting in increased levels of the transporter at the membrane and high B concentrations resulting in the transporter being targeted for degradation (Takano et al., 2005).

Plants have been shown to exhibit differences in their sensitivities to B deficiency and in the patterns of expression of BOR proteins. *A. thaliana* has previously been reported to succumb to B deficiency at levels less than 0.5 μM B, whereas rice (*Oryza sativa*) is only sensitive at levels lower than 0.2 μM B indicating that *O. sativa* is more tolerant to B deficiency (Miwa et al., 2006). In *A. thaliana*, BOR1 is expressed in the root endodermis. Whereas BOR1 from *Oryza sativa* (rice), OsBOR1, is expressed in both the endodermis/stele (for xylem loading) and in the exodermis under conditions of B deficiency. However, under conditions of B sufficiency OsBOR1 expression is only detectable in the stele (Nakagawa et al., 2007). In both species BOR1 allows passage of B though the Casparian strips thus enabling active efflux of B from the root cells to the xylem (Nakagawa et al., 2007).

BOR proteins belong to the Solute Carrier (SLC) 4 family of secondary active transporters, that includes proteins which function as either symporters or exchangers (Bai et al., 2017). Four structures of the SLC4 family are available: the human Anion Exchanger 1 (hAE1) (Arakawa et al., 2015), the acid-base transporter NBCe1 (Huynh et al., 2018), *Saccharomyces mikatae* Bor1p (Coudray et al., 2017) and *Arabidopsis thaliana* BOR1 (AtBOR1) (Thurtle-Schmidt and Stroud, 2016). The structures revealed a protomer arrangement of 7+7 transmembrane (TM) segments in an inverted repeat architecture. Each protomer is comprised of two different domains, the core and the gate, responsible of substrate transport and dimerization respectively. The SLC4 proteins are structurally related to the SLC26 family including the structurally characterised SLC26Dg (Geertsma et al., 2015; Chang et al., 2019) and BicA (Wang et al., 2019), as well as the SLC23 family including structurally characterised UapA (Alguel et al., 2016) and UraA (Yu et al., 2017). All of these proteins are dimeric and suggested to function via an elevator mechanism where substrate binds into a site formed by two half helices, TMs3 and 10, within the core domain and this domain then moves relative to the gate domain pulling the substrate across the membrane (Boudker et al., 2007; Drew and Boudker, 2016). In the case of UapA and UraA it is clear that the dimer is critical for function whereas the role of the dimer in the other proteins is less defined. Research on the *S. cerevisiae* homologue, ScBOR1p, supported the dimer as the oligomeric form of the protein and found that dimer formation was dependent upon the presence of membrane lipids, but also showed that the protein could function as a monomer (Pyle et al., 2019).

Despite progress in understanding of the structure and function of the BOR proteins, many questions remain. Here we used a combined structural and functional approach including plant phenotypes to explore the AtBOR1 and OsBOR3 proteins. Our research shows that the equivalent of disease-causing mutations from hAE1 also have highly detrimental effects on BOR protein function and plant growth. Specifically mutations G796R (Iolascon et al., 2009)) and S762R (Guizouarn et al., 2011) in hAE1 abolish anion exchange and cause stomatocytosis. The equivalent mutations in AtBOR1, A500R and S466R, respectively cause reduced boron efflux leading to boron deficiency symptoms in plants including reduced plant weight. The same results are seen for a mutant of D311, a likely proton binding residue. A low resolution cryo-EM structure of OsBOR3 provides the first glimpse of the molecular details of a rice borate transporter.

## Results

### Mutations on AtBOR1 were selected based on homology with hAE1

D311 is a possible candidate for H^+^ binding residue and mutation of the equivalent residue (D347) in *S. cerevisiae* has previously been shown to inhibit boron efflux function in a yeast based functional complementation assay (Thurtle-Schmidt and Stroud, 2016). As described above S466R (equivalent to S762R (Guizouarn et al., 2011)) and A500R (equivalent to G796R (Iolascon et al., 2009)) are equivalent to disease causing mutations in hAE1. In the context of the hAE1 structure, these mutations cannot be accommodated without modification to the overall fold (Arakawa et al., 2015). We expected P362G to be a neutral mutation with no effect on AtBOR1 transport function (see results below) and to act as a control for the generation of the plant variants.

### The AtBOR1 variants localise to the plasma membrane in yeast but exhibit varying stability in detergent based solution

All variants had similar expression levels in yeast to WT AtBOR1 (range 1.7-3 mg/L) with the exception of the A500R construct which exhibited markedly lower expression (0.7 mg/L, Supplementary Table 2). All constructs traffic to the plasma membrane in yeast (Figure 2), although unsurprisingly the level of plasma located A500R protein was lower than for the other constructs. However in the case of all the proteins there is a high level of fluorescent material clearly visible within the cells. It is likely that this is incorrectly folded material that is retained within the ER and ultimately targeted for degradation (Phillips et al., 2020).

**Figure 1.**
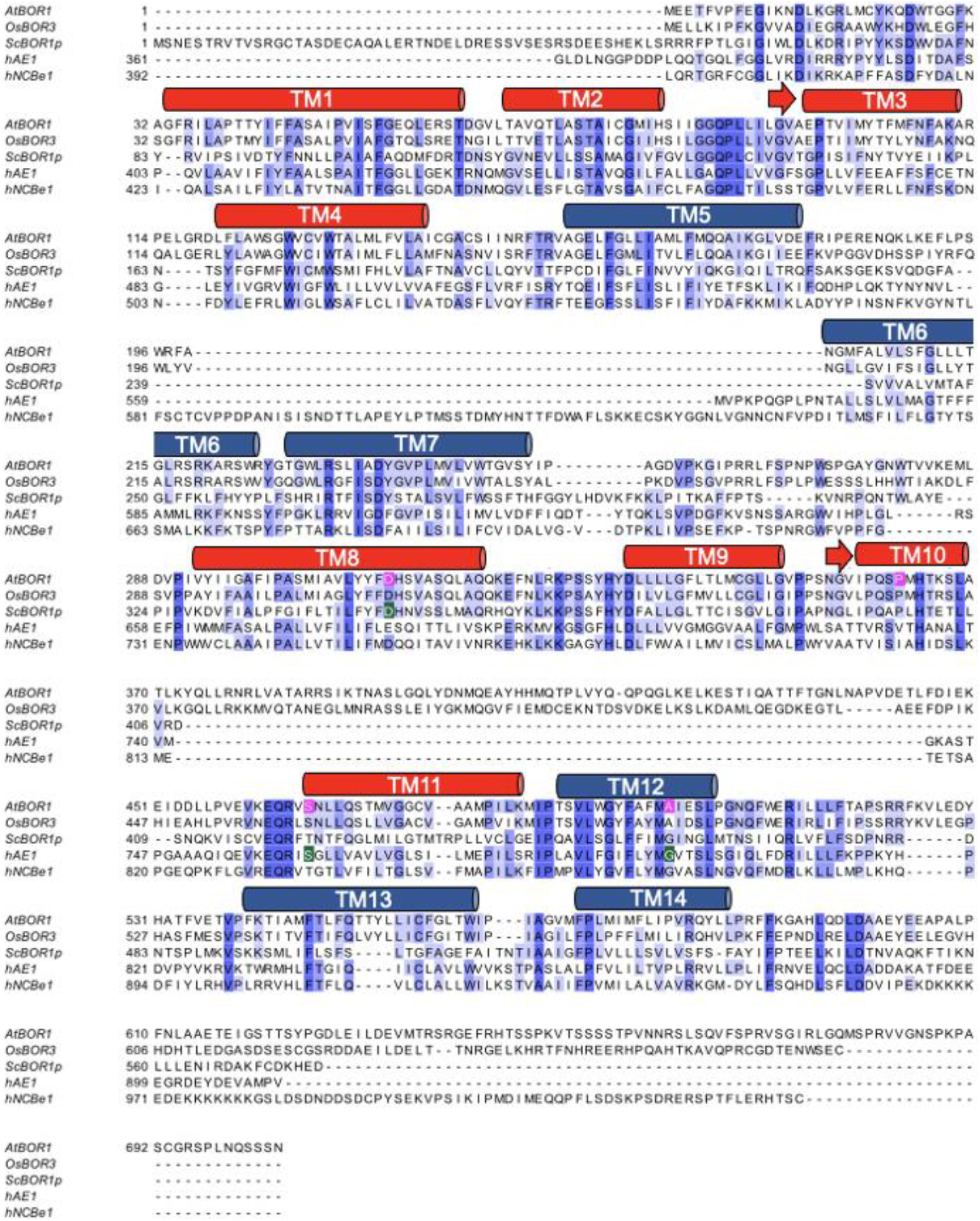
Protein sequence alignment of SLC4 family members. AtBOR1 (Acc. Code: OAP07993.1), OsBOR3 (Acc. Code: AK072421), hAE1 (Acc. Code: NP_000333.1), hNCBe1 (Acc Code:NP_003750.1). Only the TM domains of hAE1 and hNCBe1 are aligned with the full plant BOR proteins. The approximate positions of the TM domains based on the X-ray crystal structure of AtBOR1 (PDB 5L25) are indicated in the cylinders above the sequence. The TM domains are coloured with the helices comprising the core domain in red and the helices comprising the gate domain in blue. The positions of the single point mutations introduced in AtBOR1 are indicated in magenta.

**Figure 2.**
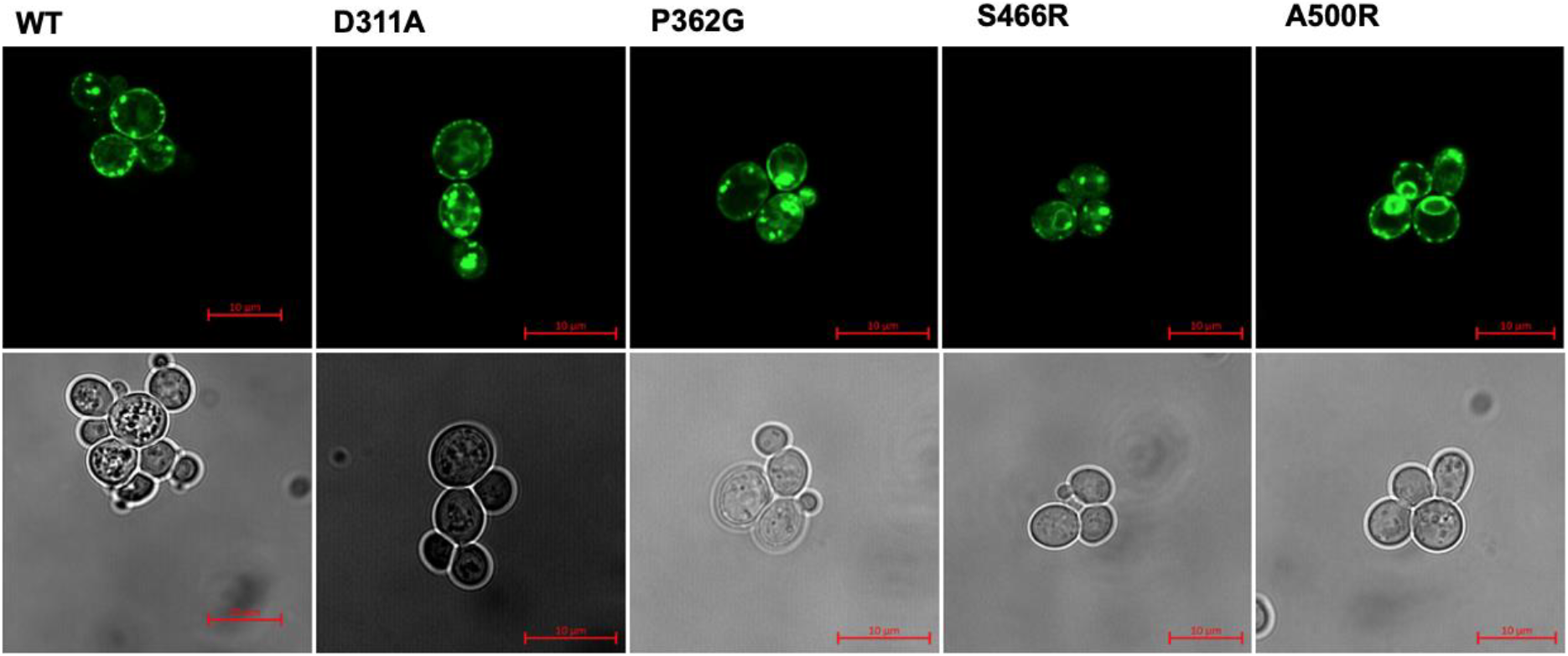
Localisation of AtBOR1-GFP variants in yeast strain FGY217. The proteins were visualised using both confocal fluorescence microscopy for detection of GFP (upper panel) and corresponding bright field images (lower panel). Data shown is representative of at least n = 2 independent experiments. Scale bar = 10 μm.

In order to determine the stability of the AtBOR1 variants we used fluorescent size exclusion chromatography (FSEC) and heated FSEC (hFSEC). All constructs solubilised into detergent (DDM)-based solution with similar solubilisation efficiencies (50-80%, **Supplementary Table 2**), although the amount of monodispersed protein differs as a result of variations in the protein expression level, particularly with respect to S466R (**Figure 3; Supplementary Table 2**). The exception was A500R where the amounts of protein extracted into detergent (29%, **Supplementary Table 2**) were too low to be detectable following separation on the SEC column. In order to further characterise the individual proteins we also submitted them to hFSEC, heating the solubilised constructs to 46ºC (higher than the apparent T_m_ for WT AtBOR1 fused with GFP; Cecchetti et al, Mansucript in preparation) for 10 mins prior to separation on the SEC column. Following heating the WT monodispersed protein peak height is substantially reduced with a concomitant increase in the size of the aggregation peak. The data revealed that the S466R was much less stable than the WT; after heating there was no detectable monodispersed protein. In contrast, the D311A and P362G constructs gave very similar results to the WT protein following heating (**Figure 3**).

**Figure 3.**
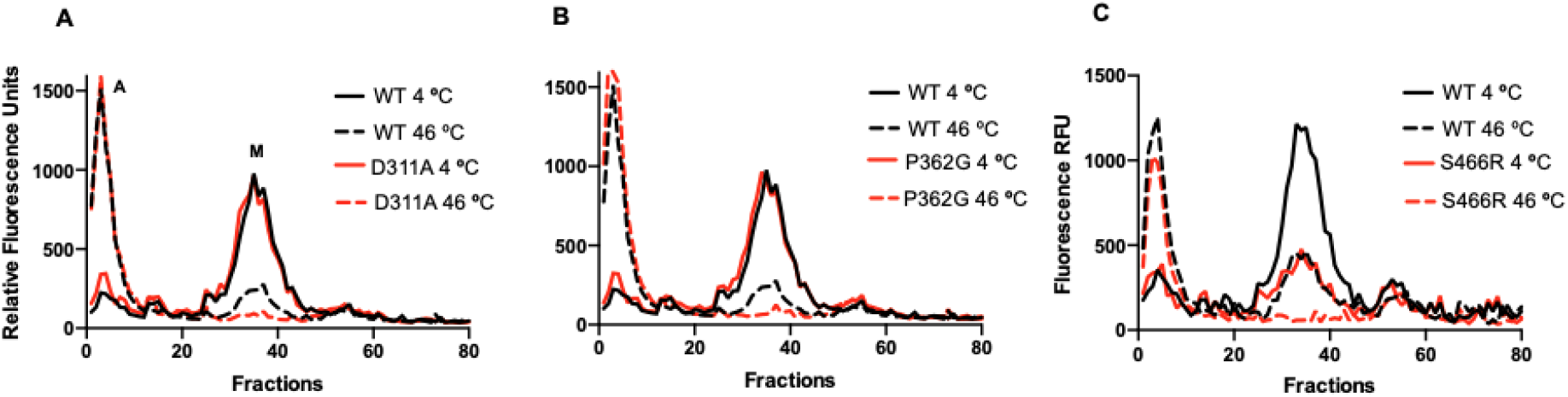
FSEC and hFSEC analysis of AtBOR1 constructs. FSEC analysis is carried out on protein solubilised at 4ºC prior to separation on the SEC column and hFSEC analysis is carried out on protein initially prepared in the same way then heated at 46ºC for 10 mins, prior to separation on the SEC column. In each case, aggregated material elutes at roughly fraction 8 with monodispersed protein eluting at roughly fraction 35, labelled A and M respectively on panel A. **A)**, **B)** and **C)** show the results obtained for D311A, P362G and S466R variants compared to WT AtBOR1 analysed at the same time, respectively. In each case the elution trace obtained for unheated sample is shown in the solid line while the elution trace obtained for heated sample is shown in the same coloured dashed line. Data is representative of at least n=2 independent experiments.

### Boron efflux was disrupted by mutations D311A, S466R and A500R

The boron efflux activity of all the constructs was assessed by expressing the constructs in a Δ*bor*1p strain. Yeast cells expressing both WT AtBOR1 and P362G exhibit reduced levels of intracellular boron compared to cells containing a vector only control (**Figure 4**). D311A in contrast exhibits a similar level of intracellular boron to the vector only control indicating no discernable boron efflux activity and in agreement with the results obtained for ScBOR1p equivalent mutant (D347; (Thurtle-Schmidt and Stroud, 2016)). Yeast cells expressing the S466R and A500R are also unable to export excess boron with similar levels of intracellular boron to the vector only control.

**Figure 4.**
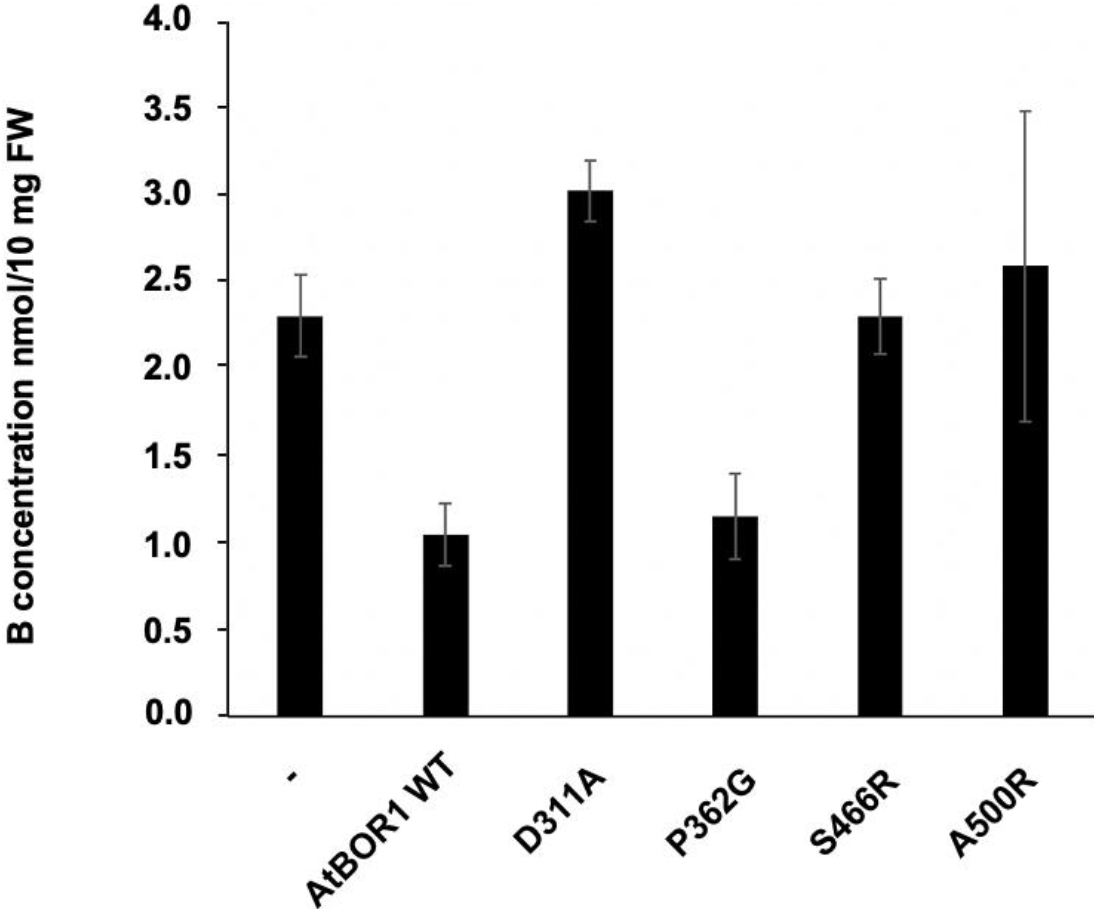
Boron efflux activity of the AtBOR1 variants in yeast. FGY217 Δ*bor1p* cells expressing each of the AtBOR1 constructs were assessed for their ability to export boron relative to control cells transformed with vector only. - = vector only control. Intracellular B content (nmol/10 mg FW) after 60 minutes incubation of cells in 1 mM boric acid. Values have been normalised to GFP concentration. Data shown is mean ± S.D. of n=3 independent experiments.

### AtBOR1 variants affect plant growth

When expressed as C-terminal GFP fusion proteins in *bor1-3*, only P362G restores shoot fresh weight and rescues B-deficiency symptoms, similar to WT BOR1 in soil-grown plants. In agreement with the results of the yeast B efflux assays D311A, S466R and A500R-expressing plants exhibit reduced shoot fresh weight and typical B-deficiency symptoms comparable to the uncomplemented *bor1-3* KO (**Figure 5**).

**Figure 5.**
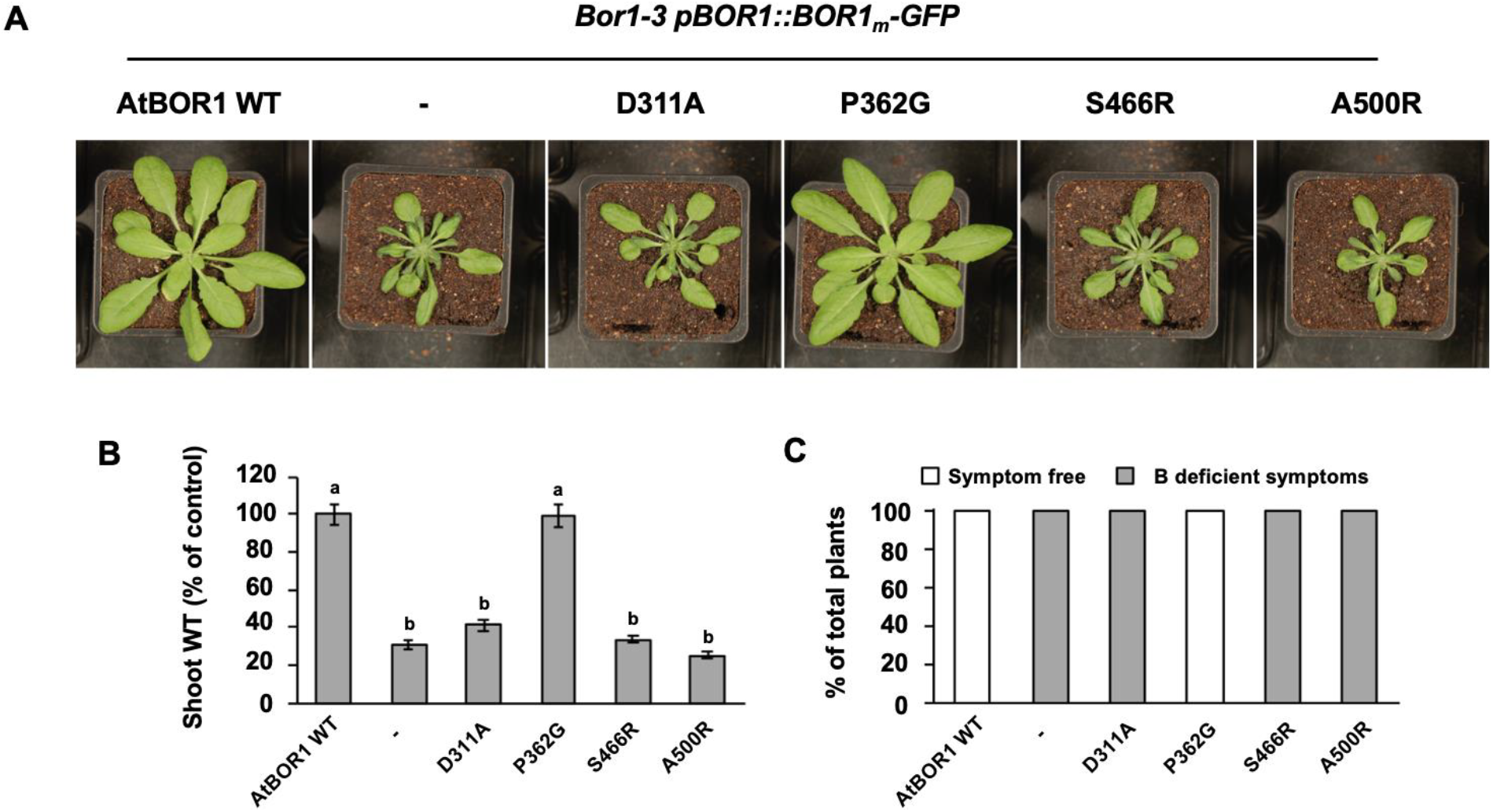
Phenotypes of different AtBOR1 variant plants grown in soil. The negative (−) control is the *bor1-3* KO plant. **A** Images of the different plants grown in soil without addition of boron. **B** Comparison of the fresh weight of the shoots of the different AtBOR1 variant plants, a, p < 0.01 compared to the negative control of the *bor1-3* KO plants, b, p <0.01 compared to WT AtBOR1 (ANOVA followed by Tukey’s test). Data is expressed relative to WT AtBOR1 and is mean ± S.E. of n ≥ 9 independent plants. **C** Overall % of plants exhibiting either no B deficiency (white) or B deficiency (grey) for each of the tested variants. Data is representative of 3 independent experiments.

### Variability of B deficiency phenotypes in planta

We observed that the B deficiency phenotype of D311A in soil grown plants appeared to be variable (possibly due to watering regime, humidity or soil batches). To better control boron concentration, we grew *A. thaliana* seedlings on agar plates supplemented with boric acid. When grown axenically on medium with low boron concentration (300 nM) seedlings of *bor1-3* plants expressing S466R and A500R exhibit reduced shoot weight similar to uncomplemented *bor1-3* plants, while P362G and WT BOR1 can restore normal shoot development under B-deficiency conditions (Figure 6). Unexpectedly, growth of seedlings expressing D311A were comparable to WR AtBOR1 at 300 nM boron (based on three independent lines). The observed shoot phenotypes for the S466R and A500R variants are a result of low B supply, since all tested mutants develop similar to control plants when grown on sufficient B levels (30 μM).

**Figure 6.**
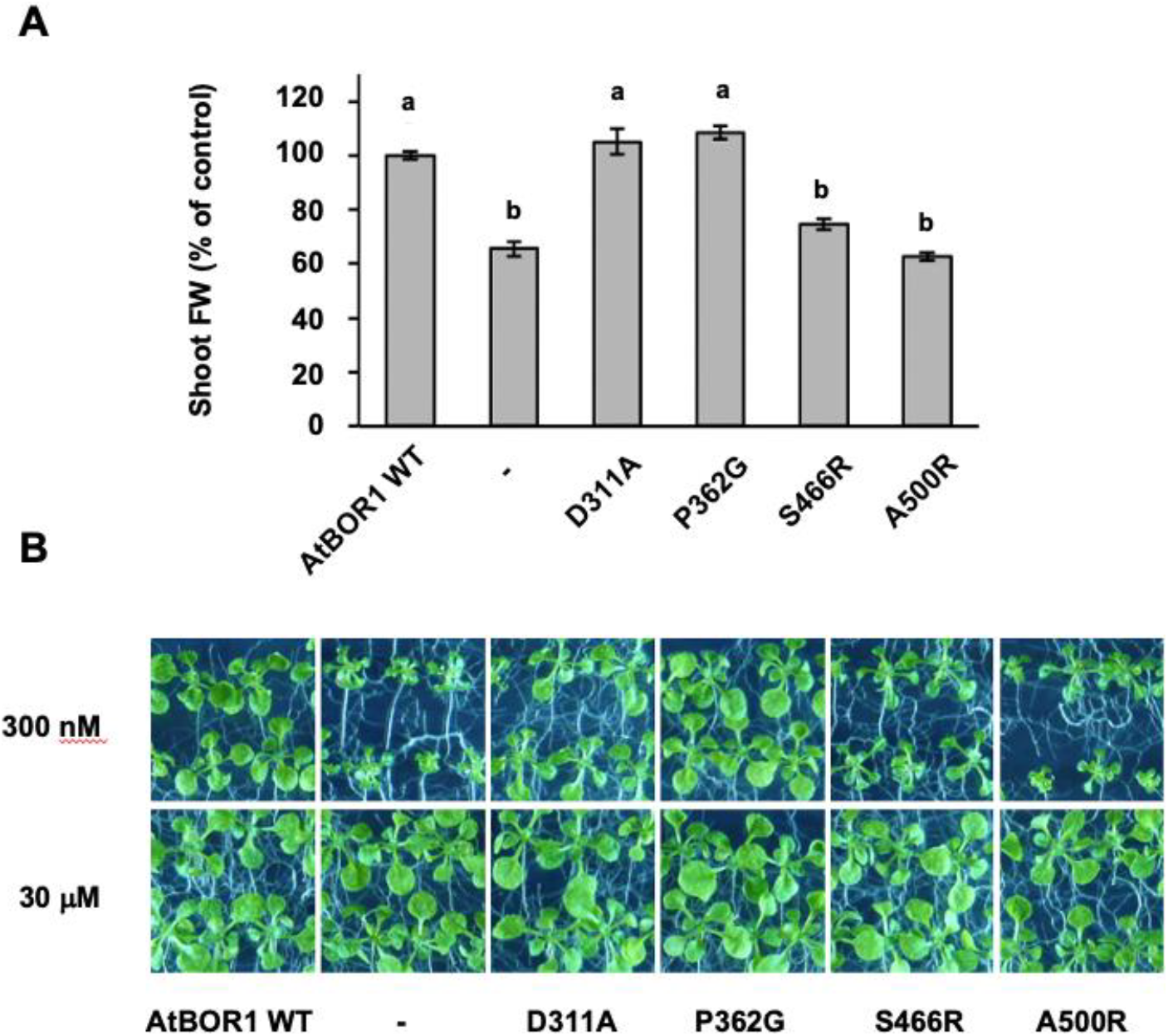
Phenotypes of AtBOR1 variant plants grown *in vitro*. **A** Comparison of the fresh weight of the different AtBOR1 variant plants grown on plates containing 300 nM boron, *a*, p < 0.01 compared to the negative control of the *bor1-3* KO plants, *b*, p <0.01 compared to WT AtBOR1 (ANOVA followed by Tukey’s test). Data is expressed relative to WT AtBOR1 and is mean ± S.E. of n = 12 plants. **B** Images of AtBOR1 variant plants grown on plants on 300 nM boron (upper panels) or 30 μM boron (lower panels). Data is representative of n = 2 independent experiments.

### OsBOR3 is a dimer, with each protomer organised into two discrete domains

There is a relatively low resolution X-ray crystallographic structure of a C-terminally truncated AtBOR1 (Thurtle-Schmidt and Stroud, 2016) in the occluded conformation (closed to both sides of the membrane). In an effort to obtain additional information about the structure and mechanism of the BOR proteins we expressed and isolated a rice homologue, OsBOR3, which shares 56% sequence identity with AtBOR1 (**Figure 1**). The protein was engineered to remove the 30 C-terminal residues and introduce a Q228T mutation to remove a naturally occurring TEV cleavage site to yield OsBOR3_Δ1-642_. Functional complementation in yeast confirmed that this engineered version of the protein is functional (**Supplementary Figure 1A**). As reported previously we have been able to obtain high quality OsBOR3_Δ1-642_ protein suitable for structural analysis (Saouros et al., 2020) (**Supplementary Figure 1B**). The protein yielded high quality negative stain and cryo-EM grids (**Supplementary Figure 1C and D**). 2D class averages were obtained of various views of the protein from a total of 116 140 particles (**Supplementary Figure 1D).** These were used to reconstruct a 3D model with a local resolution ranging from 5-6 Å **(Figure 7A)**. As expected from our previous analysis the protein is a dimer in LMNG micelles (Saouros et al., 2020). The density map for the dimer is shown in **Figure 7A** with the crystal structure of AtBOR1 fitted. We carried out separate fitting of the gate and core domain structures of outward facing hNBCe1, outward facing hAE1 and inward occluded AtBOR1 (**Figure 7B)**. As can be seen, despite the different conformational states, the domain structures of the different proteins converge on the OsBOR3 density confirming the organisation of the individual TMs into gate and core domains.

**Figure 7.**
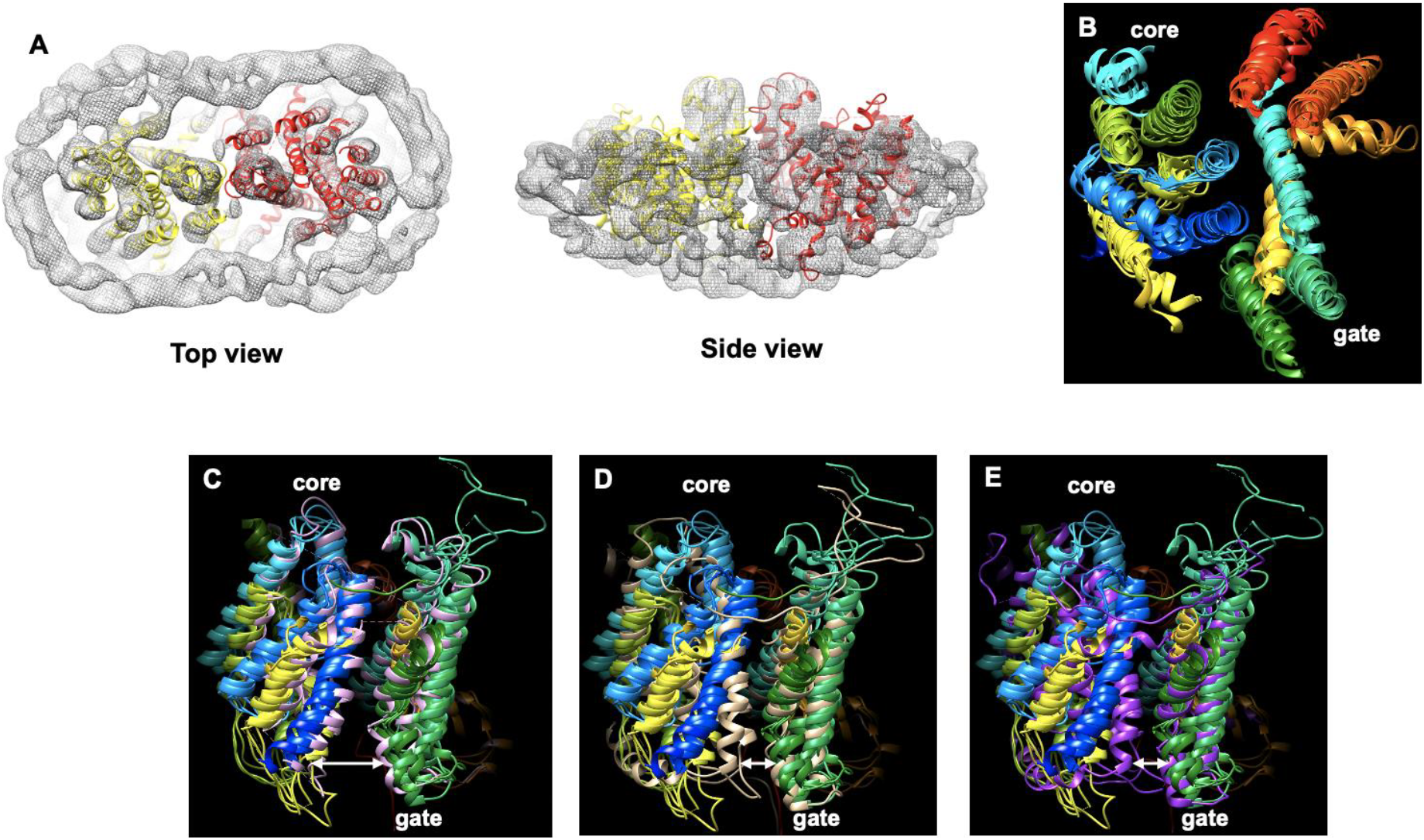
Cryo-EM structure of OsBOR3. **A)** CryoEM density map for OsBOR3 shown from the extracellular side of the membrane and the side view looking through the membrane. The full dimeric structure of AtBOR1 (individual protomers shown in yellow and red) has been fitted into the OsBOR3 EM density. The additional density around the TM domains corresponds to the LMNG detergent micelle. **B)** The respective core and gate domains of AtBOR1, NBCe1 and hAE1 fitted into the EM density of OsBOR3. A single subunit of each protein was first fitted into the density and then the automatic fitting algorithm in Chimera was used to optimise the fit of the core and gate domains separately. Each protein has been coloured in the same way with blue at the N-terminus through to red at the C-terminus. This image reveals the relative positions of the core and gate domains in the OsBOR3. Superposition of the monomeric AtBOR1 (pink **C**), hNBCe1 (beige, **D**) and hAE1 (purple, **E**) onto the composite image shown in **B**) revealing the OsBOR3 has a conformation most similar to the inward occluded AtBOR1. The images are shown from the side of the protein, looking through the membrane with the extracellular regions of the proteins should at the top. The arrows highlight the more open nature of the OsBOR3 on the intracellular side of the membrane compared to hNBCe1 and hAE1.

The core domain is less well resolved than the gate domain, probably as a result of the inherent flexibility of this region of the protein, associated with its elevator motion through the membrane. The gate domain forms the dimerization interface and as a result is much more static (**Supplementary Figure 3**).

The overall conformation of the protein after fitting the core and gate domains separately is more similar to the inward-occluded AtBOR1 (**Figure 7C**), rather than the outward-facing hNBCe1 or hAE1 (**Figure 7D, E**), indicating that OsBOR3 is in the inward facing conformation. Indeed it is possible that the OsBOR3 is in a slightly more inward open conformation than AtBOR1 although at this low resolution it is difficult to be confident of this.

## Discussion

The BOR proteins are crucial for efficient uptake and distribution of boron, as well as regulation of intracellular boron concentrations in plants. Here we used a combined approach incorporating mutagenesis in both yeast and plants, together with structural analysis in order to explore in more detail the mechanism of action of these important proteins. A series of AtBOR1 variants were generated based on i) a known loss of function mutant identified in the ScBOR1p (D311A) and ii) disease causing mutations in the related SLC4 protein, hAE1 (S466R and A500R). We also included a mutation, P362G, which had no significant effect on the expression, trafficking or function of the transporter.

The mutagenesis studies in yeast and plants were carried out using the AtBOR1 protein while the structural analysis was carried out on the OsBOR3 protein. Use of *Arabidopsis thaliana* for mutagenesis allowed us to generate plant variants on a reasonable timescale, something that would not have been possible using the OsBOR3 protein and rice plants. However, it is clearly important that we study the rice variants of these proteins given the importance of rice as a staple food for large swathes of the world’s population. In our hands, we found the C-terminally truncated OsBOR3_Δ1-642_ protein to be an extremely stable BOR variant (Saouros et al., 2020).

The AtBOR1 D311A is equivalent to the ScBOR1p D347A mutant. In the paper describing the AtBOR1 structure, the authors used the yeast ScBOR1p as a substitute for exploring the potential role of specific residues in substrate binding and transport in a functional complementation assay using a yeast Δ*bor1* strain (Thurtle-Schmidt and Stroud, 2016). They reported that transformation with AtBOR1 failed to functionally complement the ScBOR1p boron efflux activity. However in our hands we consistently saw that WT AtBOR1 was able to effectively complement ScBOR1p activity, thus allowing us to directly assess the effects of mutations in the plant transporter. In agreement with the earlier study on the ScBOR1p, here we report that mutation of AtBOR1 Asp311 to Ala results in a loss of boron efflux activity. Importantly however we also show for the first time what effect this mutant has on plant growth under boron restricted conditions. It is clear that the D311A plant variant exhibits very poor growth in soil, as poor as the *bor1-3* KO plants. Importantly, this variant expresses to a similar level to WT, traffics to the membrane and can be solubilised effectively into detergent based solution indicating that this loss of function is not due to reduced expression or the expression of incorrectly folded protein. These data confirm that this Asp residue which is conserved among BOR proteins is crucial for function. D311A is located in the TM8 which forms part of the core domain. Comparison with the structures of other SLC4 proteins, hAE1 and hNCBe1, reveals that this residue lies in close proximity to the likely substrate binding site between the two half helices, TM3 and TM10. Thus, this residue may have a role in direct substrate binding or may be the proton binding residue critical for energising the transport process. It is also possible that this residue has key roles in both substrate and proton binding.

hAE1 is by far the best characterised of the SLC4 transporters. This protein is expressed in large quantities naturally in erythrocytes and functions as a bicarbonate/Cl^−^ exchanger increasing the CO_2_ carrying capacity of the blood (Ho and Guidotti, 1975). Mutations in this protein are associated with a wide range of blood diseases (Bruce et al., 2005). These include the S762R (S466R in AtBOR1) and G796R (A500R in AtBOR1) individual mutants characterised by a loss of anion exchange activity and leakage of monovalent cations through hAE1, responsible for hereditary forms of stomatocytosis (Iolascon et al., 2009; Guizouarn et al., 2011). Mapping of these mutations onto the structure of hAE1 indicated that the S762R mutation, located at the N-terminal end of TM11, reduces the interactions between TM10 and TM11, TM11 and TM1, and TM11 with a loop connecting to TM3 (Arakawa et al., 2015). The G796R mutation located in the middle of TM12 is suggested to alter helix packing. Thus, both mutations are likely to affect the overall structural stability of the protein, although this has not been definitely demonstrated. We assessed what effects the equivalent mutations have on the expression, localisation and activity of AtBOR1. The S466R and A500R AtBOR1 variants express in yeast although both exhibit markedly lower expression than the WT transporter. Importantly, both variants do localise at the plasma membrane, suggesting that at least some of the protein is correctly folded and trafficked. FSEC and hFSEC analysis were not possible for the A500R mutant. This is most likely due to the low expression level, however we cannot rule out that low stability of this construct means that the protein does not go into detergent based solution but rather aggregates upon solubilisation. In contrast, the S466R mutant was extracted into DDM-based solution effectively, although from the hFSEC analysis is less stable than the WT protein. Boron efflux analysis revealed that both mutants exhibit a loss of transporter activity similar to the D311A mutant and this is further clearly illustrated by the poor growth of the S466R and A500R plant mutants grown in both soil and on media under boron restricted conditions. It is possible that some of the loss of function for these mutants, and particularly the A500R, is due to low expression of these constructs. In the case of the hAE1 mutants it is clear that fundamental changes within the protein abolish bicarbonate/Cl-exchange and induce cation leak (Iolascon et al., 2009; Guizouarn et al., 2011). Given the similarity between the BOR proteins and hAE1 it is likely that the mutations have similar effects at the molecular level reinforcing the suggestion that the proteins function via the same overall mechanism.

Our structure confirms findings from earlier studies on other homologous proteins that OsBOR3 is dimeric with each protomer organised into core and gate domains. In our case the core domain was slightly less well resolved compared to the gate domain, most likely as a result of the elevator motion carried out by this domain as part of the transport cycle. We did not lock our protein through the introduction of mutations or the use of a conformation specific antibody, but either of these approaches might facilitate obtaining a higher resolution structure.

In summary we have demonstrated that AtBOR1 mutations, equivalent to disease causing mutations in hAE1, result in markedly reduced plant growth phenotypes.

## Materials and Methods

### Expression, purification and cryo-EM grid preparation

The proteins were expressed and purified as previously described (Pyle et al., 2019; Saouros et al., 2020). In brief, for cryo-EM analysis a construct lacking amino acid residues 643–672 and incorporating a Q228T mutation was introduced to remove a naturally occurring TEV cleavage site (using oligomers: O_SB3_Q-T_F 5’-tctgagccaccccgtaccatacacccatgaccttgccc-3’ and O_SB3_Q-T_R 5’-gggcaaggtcatgggtgtatggtacggggtggctcaga-3’) was used. The construct, known as OsBOR3_Δ1-642_, was expressed as a fusion protein with C-terminal yeast enhanced GFP and 8 His tags and incorporating a TEV cleavage site (Drew et al., 2008). The protein was expressed in FGY217 *Saccharomyces cerevisiae* cells (Woolford et al., 2010) in a typical culture volume of 12 L -URA medium supplemented with 0.1% glucose incubated at 30ºC. Addition of galactose, to a final concentration of 2%, was used to induce protein expression once the culture reached an OD_600_ = 0.6. Protein expression was induced for 22 hours and the cells harvested by centrifugation (5000 *g* for 10 mins) followed by storage at −80ºC until further use. A 2 day protocol was used for preparation of cryo-EM grids including membrane preparation, protein isolation and grid preparation and freezing. The cells (from 4 L of culture) containing OsBOR3_Δ1-642_ were thawed and then resuspended in 50 mM Tris (pH 7.5), 1 mM EDTA, 0.6 M sorbitol) supplemented with cOmplete™ EDTA free protease inhibitor tablets (Roche). The cells were lysed using a Constant Systems Cell Disrupter and membranes harvested by differential centrifugation (10,000 *g* for 10 mins followed by 100,000 *g* for 1 hr). The membranes were immediately resuspended into solubilisation buffer (1x PBS (pH 7.5), 100 mM NaCl, 10% glycerol (v/v), 1% LMNG (w/v)) supplemented with protease inhibitor tablets (Roche), followed by incubation at 4ºC for 1 hr. Insoluble material was removed by ultracentrifugation (100,000 *g* for 1 hr) and the OsBOR3_Δ1-642_ in the soluble extract was purified using a three step process using LMNG at a concentration of 1 x CMC; His-trap, TEV cleavage followed by reverse His-trap and size exclusion chromatography. The protein was eluted from the SEC column in 20 mM Tris (pH 7.5), 150 mM NaCl, 0.75 x CMC LMNG. The single SEC fraction containing the highest concentration was used to prepare the grids directly with no further buffer exchanges or concentration steps. The protein was also analysed by Coomassie stain SDS-PAGE.

### Negative stain TEM

Protein preparations were screened by negative-stain EM to assess for monodispersity and overall quality of the protein preparation prior to cryo-EM analyses. For negative strain EM analysis, 0.01 mg/ml OsBOR3_Δ1-642_ in a total volume of 3 μl was applied onto freshly glow-discharged carbon coated 400 mesh grids (Agar scientific), excess solution removed by blotting from the side and stained with 2% uranyl acetate. Negative stain EM micrographs were recorded on a Tecnai T12 microscope at 120 kV using an FEI 2K eagle camera.

### CryoEM grid preparation and data collection

300 mesh Cu Quantifoil R2/2 grids (Agar Scientific) were glow discharged for 90 s prior sample deposition. 3 μL OsBOR3 (1 mg.mL^−1^) was applied to the grid, blotted and then plunged into liquid ethane using a Vitribot mark IV (Thermo Fisher Scientific) at 4 ºC, 100% humidity, 3 s blot time and −2 blot force. Plunge frozen grids were stored under liquid nitrogen until use.

EPU automated imaging software (Thermo Fisher Scientific) was used to collect electron micrograph movies on a 300 keV Titan Krios (Thermo Fisher Scientific) fitted with a K3 direct detection camera (Gatan) in super resolution mode. All collection parameters are shown in Supplementary Table 3.

### CryoEM data processing and density reconstruction

The collected micrograph movie frames were aligned using MotionCor2 (Zheng et al., 2017) and subsequently the CTF parameters were estimated with CTFFind4 (Rohou et al, 2015). All the micrographs were then assessed by eye for quality using the FFT spectra to discard micrographs with excessive image drift or crystalline ice. Unless otherwise stated all the following data processing was carried out in RELION (Zivanov et al., 2018). Particles were picked manually and with the RELION auto-picking function and then sorted by their similarity to the reference 2D classes (Z-score). Several rounds of 2D classification were used to select a range of orientations before generating an initial model low pass filtered to 15 Å. A three class 3D classification was performed with the initial model to select a homogenous particle set for refinement. The class with the highest particle occupancy and resolution was taken forward for refinement. 3D refinement with C2 symmetry imposed, followed by post-processing yielded a reconstructed density of between 5 and 6 Å estimated global resolution with a B-factor of −220 Å^2^. Details of the mask used are available from the Electron Microscopy DataBank (EMDB) under the accession code EMDB-11996. All maps were visualised in 3D using UCSF Chimera (Pettersen et al., 2004).

### Mutagenesis

AtBOR1 point mutants were created with the QuikChange Lightning Site-DirectedMutagenesis Kit (Agilent Technologies), using as template DNA, either wild-type or truncated AtBOR1 in the pDDGFP2 vector. The oligonucleotides used are shown in Supplementary Table 1.

### Functional complementation analysis

pDDGFP2 vectors containing the genes encoding WT AtBOR1, WT OsBOR3 and OsBOR3_Δ1-642_ were transformed into a *S. cerevisiae* Δ*bor*1p strain for functional complementation analysis as previously described (Pyle et al., 2019). As a negative control empty pDDGFP2 vector was also transformed into the *S. cerevisiae* Δ*bor*1p strain. Transformants were incubated in 10 mL -URA with 2 % glucose at 30 °C and 300 rpm shaking overnight. A 5-fold serial dilution was carried out with cells starting at OD_600_ 0.5 in -URA media. 10 μL of each dilution was spotted on -URA plates with 2 % galactose and supplemented with the following concentrations of boric acid: 0, 2.5, 5, 7.5 mM. Plates were incubated for 30 °C for 5 or 7 days and imaged using a ChemiDoc MP Imaging System (Bio-Rad).

### FSEC and hFSEC

AtBOR1 constructs were expressed as described above and membranes containing the different AtBOR1 proteins were diluted in solubilisation buffer (PBS (pH 7.4), 1% DDM, supplemented with 1 tablet of protease inhibitor) to give a final total protein concentration of 50 μg/ml and incubated with gentle rocking at 4 °C for 1 hour. Insoluble material was removed by centrifugation at 14 000 g for 1 hour at 4 °C. The solubilisation efficiency of each construct was calculated as the proportion of the initial GFP fluorescence that remained after solubilisation. 500 μl supernatant was injected onto a Superose 6 10/300 column equilibrated with 20 mM Tris (pH 7.5), 150 mM NaCl and 0.03% DDM. The elution was collected, from 6.4 ml elution volume (void), in 200 μl fractions in a clear bottomed 96-well plate. The GFP fluorescence of each fraction was measured using a SpectraMax M2e with an excitation wavelength of 488 nm and an emission wavelength of 512 nm. For heated FSEC experiments, the solubilised protein was incubated at 46°C for 10 minutes and any insoluble aggregates removed by a 10 minute centrifugation prior to loading onto a Superose 6 10/300 column. The samples were collected and analysed as described above.

### Plant genotypes

Using *Agrobacterium tumefaciens*-mediated transformation, the pBOR1::BOR1-GFP constructs containing the respective point mutations were introduced into the *bor1-3 Arabidopsis thaliana* background, which carries a T-DNA insertion in the AtBOR1 gene (SALK_037312, (Miwa et al., 2013)). To select for plants carrying the desired transgene, T1 seeds were grown on half strength Murashige and Skoog (MS) medium containing 50 mg/L kanamycin. The presence of the respective BOR1 mutation in each individual line was verified by PCR on genomic DNA (primers 5’-ATGCTTGATGTTCCAATCGTC-3’ and 5’ AGCTCCTCGCCCTTGCTCAC-3’) followed by sequencing. The resulting T2 seeds were analysed for their segregation on kanamycin to avoid lines containing multiple insertions of the construct. Several independent insertion lines were isolated per construct to minimize the risk of positional effects. The T2 seeds were used for phenotyping experiments.

### Plant growth conditions

For analysis of shoot fresh weight of seedlings grown in axenic culture, plants were grown in square petri dishes (12 × 12 cm) for 3 weeks. Seeds were surface-sterilized and stratified for 2-3 days at 4°C. Half strength MS basal medium (corresponding to Sigma M5524) was prepared by adding all ingredients except boric acid. The MS medium contained 5 g/L sucrose and 0.5 g/L MES (2-(N-morpholino)ethanesulfonic acid) and was adjusted to pH 5.7 with KOH. To remove residual boric acid the medium was treated overnight with B chelator Amberlite IRA-743 (3 g/L, Sigma). Before autoclaving, boric acid was added to the medium to a concentration of 30 μM (sufficient B) or 300 nM (low B). The medium was solidified using 4.5 g/L Gelrite (Duchefa). The seeds were germinated vertically on medium containing 50 mg/L kanamycin to select T2 seedlings that carry the desired transgene. When 5 days old, resistant seedlings were transferred to fresh medium with either low or sufficient boron but without kanamycin and grown horizontally thereafter. When the plants were 3 weeks old the shoots were separated from the root system and weighed. Plates were kept in a growth chamber set to 22°C day / 20°C night temperature under 12 h of light.

To determine plant phenotypes in soil culture, seedlings were initially grown vertically on half strength MS medium containing 30 μM boron and 50 mg/L kanamycin as described above. After 1 week, 10-12 resistant seedlings per genotype were transferred to soil into individual pots (5 cm). After transfer to soil, the plants were moved to an Aralab growth room set to 22 °C day / 20 °C night temperature under 12 h of light at 60 % relative humidity. The plants were watered moderately and the rosette fresh weight was analysed at 5 weeks of age.

### Boron transport assay

Synthetic genes of AtBOR1 mutants in pDONR221 were synthesized by Genescript (https://www.genscript.com/). The synthetic genes were subcloned into yeast expression vector pDDGFP2 and transformed in to FGY217 (Δbor1) yeast strain as described by (Pyle et al., 2019). Boron efflux assay in yeast cells was performed as described in Takano et al. (Takano et al., 2002) with some modifications. Briefly, yeast cells harvested at mid-exponential phase were transferred to the synthetic medium (pH 5.5) containing 1 mM of Boric acid and incubated for 1 hr. After 1 hr cells were washed with ice cold distilled water and harvested by centrifugation at 3000 g for 2 min. To release the intracellular boron the harvested cells were boiled at 90 °C for 30 min and the supernatant was used to measure the boron content by using the protocol as described by (Mohan and Jones, 2018).

### Confocal microscopy

Yeast cells expressing AtBOR1-GFP were harvested at mid-log phase (OD600 ~ 0.6). 100μl of these cells were aliquoted in microplate and absolute fluorescence was quantified using Fluorescence Microplate reader with excitation max 488 nm and emission max 509 nm. To assess the localization of AtBOR1-GFP, yeast cells expressing the AtBOR1 constructs were harvested at mid-log phase and analysed using Leica SP5 confocal fluorescence microscope with the excitation and detection wavelengths of 488 nm and 500–540nm, respectively.

### Deposition information

The final cryoEM density map of OsBOR3 has been deposited to the Electron Microscopy DataBank (EMDB) under the accession code EMD-11996.

## Acknowledgments

This work was also supported by Biotechnology and Biological Sciences Research Council (BBSRC) grant BB/N016467/1 awarded to BB, AC and AJ. This project also received funding from the European Union’s Horizon 2020 research and innovation programme, RAMP-ITN: Rationalising Membrane Protein Crystallisation Innovative Training Network, under the Marie Sklodowska-Curie grant agreement No 722687 (CC). The authors wish to thank Nora Cronin, facility manager of LonCEM at the Francis Crick Institute for assistance with data collection. The Electron Microscopy Facility, Centre for Structural Biology, Imperial College London was used for grid preparation, optimisation and initial Cryo-EM screening.

## References

Alguel Y, Amillis S, Leung J, Lambrinidis G, Capaldi S, Scull NJ, Craven G, Iwata S, Armstrong A, Mikros E, et al (2016) Structure of eukaryotic purine/H(+) symporter UapA suggests a role for homodimerization in transport activity. Nat Commun 7: 11336–9

Arakawa T, Kobayashi-Yurugi T, Alguel Y, Iwanari H, Hatae H, Iwata M, Abe Y, Hino T, Ikeda-Suno C, Kuma H, et al (2015) Crystal structure of the anion exchanger domain of human erythrocyte band 3. Science 350: 680–684

Bai X, Moraes TF, Reithmeier RAF (2017) Structural biology of solute carrier (SLC) membrane transport proteins. Mol Membr Biol 34: 1–32

Boudker O, Ryan RM, Yernool D, Shimamoto K, Gouaux E (2007) Coupling substrate and ion binding to extracellular gate of a sodium-dependent aspartate transporter. Nature 445: 387–393

Bruce LJ, Robinson HC, Guizouarn H, Borgese F, Harrison P, King M-J, Goede JS, Coles SE, Gore DM, Lutz HU, et al (2005) Monovalent cation leaks in human red cells caused by single amino-acid substitutions in the transport domain of the band 3 chloride-bicarbonate exchanger, AE1. Nat Genet 37: 1258–1263

Camacho-Cristóbal JJ, Martín-Rejano EM, Herrera-Rodríguez MB, Navarro-Gochicoa MT, Rexach J, González-Fontes A (2015) Boron deficiency inhibits root cell elongation via an ethylene/auxin/ROS-dependent pathway in Arabidopsis seedlings. Journal of Experimental Botany 66: 3831–3840

Chang Y-N, Jaumann EA, Reichel K, Hartmann J, Oliver D, Hummer G, Joseph B, Geertsma ER (2019) Structural basis for functional interactions in dimers of SLC26 transporters. Nat Commun 10: 1–10

Chormova D, Fry SC (2015) Boron bridging of rhamnogalacturonan-II is promoted in vitroby cationic chaperones, including polyhistidine and wall glycoproteins. New Phytol 209: 241–251

Coudray N, L Seyler S, Lasala R, Zhang Z, Clark KM, Dumont ME, Rohou A, Beckstein O, Stokes DL (2017) Structure of the SLC4 transporter Bor1p in an inward-facing conformation. Protein Sci 26: 130–145

Dell B, Huang L (1997) Physiological response of plants to low boron - Springer. Plant and Soil 193: 103–120

Drew D, Boudker O (2016) Shared Molecular Mechanisms of Membrane Transporters. Annu Rev Biochem 85: 543–572

Drew D, Newstead S, Sonoda Y, Sonoda Y, Kim H, Kim H, Heijne von G, Iwata S (2008) GFP-based optimization scheme for the overexpression and purification of eukaryotic membrane proteins in Saccharomyces cerevisiae. Nat Protoc 3: 784–798

Geertsma ER, Chang Y-N, Shaik FR, Neldner Y, Pardon E, Steyaert J, Dutzler R (2015) Structure of a prokaryotic fumarate transporter reveals the architecture of the SLC26 family. Nature Structural & Molecular Biology 22: 803–808

Goldbach HE, Yu Q, Wingender R, Schulz M, Wimmer M, Findeklee P, Baluska F (2001) Rapid response reactions of roots to boron deprivation. Journal of Plant Nutrition and Soil Science 164: 173–181

Guizouarn H, Borgese F, Gabillat N, Harrison P, Goede JS, McMahon C, Stewart GW, Bruce LJ (2011) South-east Asian ovalocytosis and the cryohydrocytosis form of hereditary stomatocytosis show virtually indistinguishable cation permeability defects. Br J Haematol 152: 655–664

Ho MK, Guidotti G (1975) A Membrane Protein from Human Erythrocytes involved in Anion Exchange. Journal of Biological Chemistry 250: 675–683

Hrmova M, Gilliham M, Tyerman SD (2020) Plant transporters involved in combating boron toxicity: beyond 3D structures. Biochem Soc Trans 37: 629–14

Hu H, Brown PH (1994) Localization of Boron in Cell Walls of Squash and Tobacco and Its Association with Pectin (Evidence for a Structural Role of Boron in the Cell Wall). PLANT PHYSIOLOGY 105: 681–689

Huynh KW, Jiang J, Abuladze N, Tsirulnikov K, Kao L, Shao X, Newman D, Azimov R, Pushkin A, Zhou ZH, et al (2018) CryoEM structure of the human SLC4A4 sodium-coupled acid-base transporter NBCe1. Nat Commun 9: 900–9

Iolascon A, De Falco L, Borgese F, Esposito MR, Avvisati RA, Izzo P, Piscopo C, Guizouarn H, Biondani A, Pantaleo A, et al (2009) A novel erythroid anion exchange variant (Gly796Arg) of hereditary stomatocytosis associated with dyserythropoiesis. Haematologica 94: 1049–1059

Jennings ML, Howren TR, Cui J, Winters M, Hannigan R (2007) Transport and regulatory characteristics of the yeast bicarbonate transporter homolog Bor1p. AJP: Cell Physiology 293: C468–C476

Li C, Pfeffer H, Dannel F, Romheld V, Bangerth F (2001) Effects of boron starvation on boron compartmentation, and possibly hormone-mediated elongation growth and apical dominance of pea (Pisum sativum) plants. Physiol Plant 111: 212–219

Miwa K, Fujiwara T (2010) Boron transport in plants: co-ordinated regulation of transporters. Annals of Botany 105: 1103–1108

Miwa K, Takano J, Fujiwara T (2006) Improvement of seed yields under boron-limiting conditions through overexpression of BOR1, a boron transporter for xylem loading, in Arabidopsis thaliana. The Plant Journal 46: 1084–1091

Miwa K, Takano J, Omori H, Seki M, Shinozaki K, Fujiwara T (2007) Plants tolerant of high boron levels. Science 318: 1417–1417

Miwa K, Wakuta S, Takada S, Ide K, Takano J, Naito S, Omori H, Matsunaga T, Fujiwara T (2013) Roles of BOR2, a boron exporter, in cross linking of rhamnogalacturonan II and root elongation under boron limitation in Arabidopsis. PLANT PHYSIOLOGY 163: 1699–1709

Mohan TC, Jones AME. (2018) Determination of Boron Content Using a Simple and Rapid Miniaturized Curcumin Assay. Bio Protoc. 8: e2703.

Nakagawa Y, Nakagawa Y, Hanaoka H, Hanaoka H, Kobayashi M, Miyoshi K, Miyoshi K, Miwa K, Fujiwara T (2007) Cell-Type Specificity of the Expression of Os BOR1, a Rice Efflux Boron Transporter Gene, Is Regulated in Response to Boron Availability for Efficient Boron Uptake and Xylem Loading. THE PLANT CELL ONLINE 19: 2624–2635

Pallotta M, Schnurbusch T, Hayes J, Hay A, Baumann U, Paull J, Langridge P, Sutton T (2014) Molecular basis of adaptation to high soil boron in wheat landraces and elite cultivars. Nature 514: 88–91

Pettersen EF, Goddard TD, Huang CC, Couch GS, Greenblatt DM, Meng EC, Ferrin TE (2004) UCSF Chimera--a visualization system for exploratory research and analysis. J Comput Chem 25: 1605–1612

Phillips BP, Gomez-Navarro N, Miller EA (2020) Protein quality control in the endoplasmic reticulum. Current Opinion in Cell Biology 65: 96–102

Pyle E, Guo C, Hofmann T, Schmidt C, Ribiero O, Politis A, Byrne B (2019) Protein-Lipid Interactions Stabilize the Oligomeric State of BOR1p from Saccharomyces cerevisiae. Anal Chem 91: 13071–13079

Rohou A, Grigorieff N (2015). CTFFIND4: Fast and accurate defocus estimation from electron micrographs. J Struct Biol 192: 216–221

Saouros S, Cecchetti C, Jones A, Cameron AD, Byrne B (2020) Strategies for successful isolation of a eukaryotic transporter. Protein Expr Purif 166: 105522

Shorrocks VM (1997) The occurrence and correction of boron deficiency. Plant and Soil 198: 121–148

Sutton T, Baumann U, Hayes J, Collins NC, Shi B-J, Schnurbusch T, Hay A, Mayo G, Pallotta M, Tester M, et al (2007) Boron-toxicity tolerance in barley arising from efflux transporter amplification. Science 318: 1446–1449

Takano J, Miwa K, Yuan L, Wirén von N, Fujiwara T (2005) Endocytosis and degradation of BOR1, a boron transporter of Arabidopsis thaliana, regulated by boron availability. Proceedings of the National Academy of Sciences 102: 12276–12281

Takano J, Noguchi K, Yasumori M, Kobayashi M, Gajdos Z, Miwa K, Hayashi H, Yoneyama T, Fujiwara T (2002) Arabidopsis boron transporter for xylem loading. Nature 420: 337–340

Takano J, Wada M, Ludewig U, Schaaf G, Wirén von N, Fujiwara T (2006) The Arabidopsis major intrinsic protein NIP5;1 is essential for efficient boron uptake and plant development under boron limitation. Plant Cell 18: 1498–1509

Tanaka M, Fujiwara T (2008) Physiological roles and transport mechanisms of boron: perspectives from plants. Pflugers Arch 456: 671–677

Thurtle-Schmidt BH, Stroud RM (2016) Structure of Bor1 supports an elevator transport mechanism for SLC4 anion exchangers. Proc Natl Acad Sci USA 113: 10542–10546

Uragachi S, Kaya Y, Hanaoka H, Fujiwara T (2014) Generation of boron-deficiency-tolerant tomato by overexpressing an Arabidopsis thaliana borate transporter AtBOR1. Front Plant Sci 5: 1–7

Wang C, Sun B, Zhang X, Huang X, Zhang M, Guo H, Chen X, Huang F, Chen T, Mi H, et al (2019) Structural mechanism of the active bicarbonate transporter from cyanobacteria. Nat Plants 5: 1184–1193

Woolford CA, Daniels LB, Park FJ, Jones EW, Van Arsdell JN, Innis MA (2010) The PEP4 gene encodes an aspartyl protease implicated in the posttranslational regulation of Saccharomyces cerevisiae vacuolar hydrolases. Mol Cell Biol 6: 2500–2510

Yu X, Yang G, Yan C, Baylon JL, Jiang J, Fan H, Lu G, Hasegawa K, Okumura H, Wang T, et al (2017) Dimeric structure of the uracil:proton symporter UraA provides mechanistic insights into the SLC4/23/26 transporters. Cell Res 27: 1020–1033

Zivanov J, Nakane T, Forsberg BO, Kimanius D, JH HW, Lindahl E, Scheres SHW (2018) New tools for automated high-resolution cryo-EM structure determination in RELION-3. Elife 7: e42166

Zheng, S. Q., Palovcak, E., Armache, J.-P., Verba, K. A., Cheng, Y., & Agard, D. A. (2017). MotionCor2: anisotropic correction of beam-induced motion for improved cryo-electron microscopy. Nature Methods 14: 1–2.

